# Induced aneuploidy disrupts MCF10A acini formation and CCND1 expression

**DOI:** 10.1101/2019.12.15.876763

**Authors:** Marcel Waschow, Qi Wang, Paul Saary, Corinna Klein, Sabine Aschenbrenner, Katharina Jechow, Lorenz Maier, Stephan Tirier, Brigitte Schoell, Ilse Chudoba, Christian Dietz, Daniel Dreidax, Anna Jauch, Martin Sprick, Carl Herrmann, Roland Eils, Christian Conrad

**Affiliations:** Division of Theoretical Bioinformatics, German Cancer Research Center (DKFZ), Heidelberg, Baden-Wuerttemberg, 69120, Germany; Division of Stem Cells and Cancer, German Cancer Research Center (DKFZ), Heidelberg, Baden-Wuerttemberg, 69120, Germany; Institute of Human Genetics, University of Heidelberg, Heidelberg, Baden-Wuerttemberg, 69120, Germany; MetaSystems, Altlussheim, Baden-Wuerttemberg, 68804, Germany; Bioinformatics and Information Mining, University of Konstanz, Konstanz, Baden-Wuerttemberg, 78464, Germany; Functional and Structural Genomics, German Cancer Research Center (DKFZ), Heidelberg, Baden-Wuerttemberg, 69120, Germany; Center for Digital Health, Berlin Institute of Health and Charité - Universitätsmedizin Berlin, 10117, Berlin, Germany

**Keywords:** 3D cell culture, high-content screening, aneuploidy, chromatin interaction

## Abstract

Abnormal karyotypes, namely aneuploidy, can be detected in nearly all cancer entities at different grades. The impact of these altering mutations on epigenetic regulation, especially on promoter-enhancer interactions are not well understood. Here, we applied a 3D model of MCF10A cells in a high-content screen to measure induced aneuploidy by RNA interference of 82 mitotic genes associated with aneuploidy and breast cancer. Perturbation of ESPL1 and TOP2A expression led to increased mitotic instability and subsequent aneuploidy and polylobed nuclei. During acinus formation these polylobed cells disrupted proper acinus rotation inhibiting the development of a hollow lumen and a polarized outer cell layer. Further, gene expression profiling identified upregulated CCND1 among other breast cancer related genes. We show that acquisition of aneuploidy affects the morphogenesis of MCF10A acini and expression of cancer relevant genes. By conducting 4C chromosome capturing experiments we linked the alteration of interactions of the promoter region to CCND1 upregulation.

## INTRODUCTION

Since Theodor Boveri’s discovery in the beginning of the last century that chromosomes can be unevenly divided between the two daughter cells (Boveri, 2008), decades followed when DNA mutations dominated functional cancer research. Today, we know that unbalanced chromosome sets can be found in nearly all tumor entities, at all variations. Such aneuploidies alter gene expression and can give rise to additional malignant features (Sotillo et al., 2010). Nonetheless, a connection of aneuploidy and gene expression changes of unmutated genes is not straight-forward, and barely studied. Differential gene expression can be caused on the genetic, epigenetic and translational level. While breast cancer, the most common form of cancer in women today (Bray et al., 2018), is often associated with aneuploidy (Cimini, 2008), experimental data as well as theoretical models suggest that acquisition of aneuploidy might be an early event in tumorigenesis (Nowak et al., 2002, Wang et al., 2014, Gao et al., 2016). This was described for triple-negative breast cancer where potential precursors exhibit karyotypes similar to advanced stages (Ottesen et al., 1995; Wang et al., 2014). The strong relationship between aneuploidy and breast cancer was further established in a study that could reliably stratify breast cancer stages using a gene signature that contained genes whose deregulation correlates with aneuploidy (Carter et al., 2006).

To investigate the impact of early aneuploidy on the morphogenesis of healthy acini we applied RNA interference of breast cancer as well as aneuploidy associated genes on Matrigel cultured MCF10A cells. MCF10A is a well-studied cell line, which shows a triple negative receptor status. Additionally, MCF10A cells can be assayed on Matrigel to form acini structures, which more closely resemble physiologic profiles of the mammary gland, when compared to traditional monolayer cell cultures (Debnath et al., 2003). The capability of forming healthy, round acini makes it a suitable model to study the effects of early aneuploidy on acinus morphogenesis in context to cancer initiation. The further progressed cell line MCF10CA (Santner et al., 2001) displays malformed tumor spheroids when cultured on Matrigel expressing breast cancer signatures and EMT markers and serves as a cancer model for MCF10A cells (So et al., 2012).

Chromosome conformation capture assays allow the analysis of chromatin interactions (Dekker et al., 2002, Simonis et al., 2006, Lieberman-Aiden et al., 2009). These interactions occur within and between chromosomes (Miele & Dekker, 2009) and are tightly regulated. The effects of chromosome interactions on gene expression are exerted over transcription factories (Osborne et al., 2004, Cook, 2010). Transcription factories facilitate the extensive interplay of all chromosomes by forming active islands of transcription (Cook & Marenduzzo, 2018). Although aneuploidy is a central property of cancer and chromosome capture technologies have been increasingly employed, the relationship of aneuploidy, chromosome conformation, and gene regulation has been poorly studied.

As an example, CCND1, which codes for a key regulator of the cell cycle (Otto & Sicinski, 2017), shows a heterogeneous relation of copy number variations (CNV) and corresponding expression. Cancer genome data available at the COSMIC database (Tate et al., 2018) suggests no correlation of CNV and expression for CCND1 in invasive breast cancer, while others report a correlation (Geiger et al., 2010, Lundberg et al., 2019). To explore *de novo* how aneuploidy affects healthy acinus morphogenesis, we first observed acinus development under the influence of siRNA-induced aneuploidy in a 3D culture based high-content screen. We found several morphology-disrupting gene candidates, and then conducted chromosome capture (4C) studies of the CCND1 promoter as viewpoint.

## RESULTS

### Identification of candidates altering the morphogenesis of MCF10A spheroids

In a high-content screen we evaluated a set of 82 genes that were previously reported to be associated with breast cancer as well as aneuploidy (Figure 1a and Supplementary Table 1). The effects of the knock-down on the morphogenesis of MCF10A acini were measured at three different time points by fluorescence confocal microscopy. Nine parameters including spheroid size, hollowness, and polarization were used to characterize alterations in morphogenesis. Principal component analysis (PCA) and subsequent k-means clustering indicated the most disrupted phenotypes (highlighted in red, purple, green and blue colors, Fig. 1b). Within these groups we selected *DIAPH3*, *ESPL1*, *H2AFX*, *SETDB2* and *TOP2A* (*siRNA italic*) for further analysis based on prevalence of viable but abnormally formed acini. From these we could identify *ESPL1* and *TOP2A* to induce the highest degree of aneuploidy by metaphase spreads and 24 multi-color FISH (Figure 1c, d).

**Figure 1:**
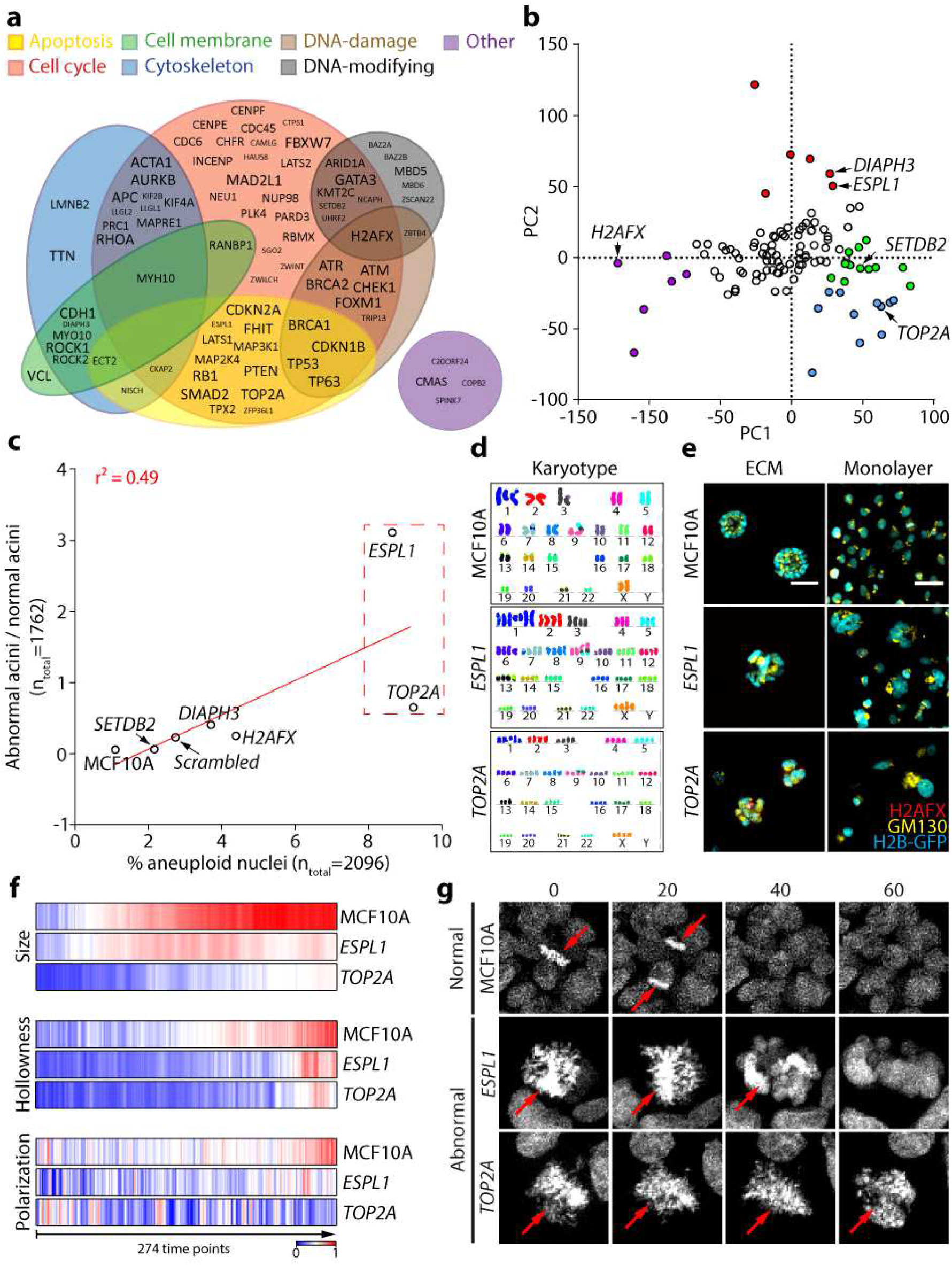
MCF10A cells form abnormal acini after induction of aneuploidy. a) Screening candidates categorized into 7 “GO-process” defined groups. The character size of the names correlates with the referenced number of publications on PubMed (big: >200 publications, middle: between 200 and 50 publications, small: < 50 publications; August 2016). b) Identification of the most disrupted siRNA-induced phenotypes by PCA and subsequent k-means clustering highlighted in red, purple, green, and blue. c) Correlation of aneuploid nuclei within acini identified by metaphase spreads (x-axis) and the ratio of morphologically abnormal and normal acini (y-axis). d) 24 multicolor FISH preparation of *ESPL1* and *TOP2A* karyotypes in comparison to normal MCF10A karyotypes. e) Depiction of the morphological phenotypes of MCF10A, ESPL1 and TOP2A acini (ECM) and monolayer cells. Red: H2AFX (DNA-damage), yellow: GM130 (Golgi), cyan: H2B-GFP (nuclei). The bar represents a length of 50 μm. f) Relative quantification of the parameters size, hollowness and polarization over 274 timepoints comparing MCF10A with *ESPL1* and *TOP2A*. g) Time-lapse images illustrating the time frame in minutes of dividing cells for MCF10A in comparison to *ESPL1* and *TOP2A*.

Cells with an aneuploid/polyploid karyotype show a characteristic irregular nucleus shape in both 3D and monolayer culture (polylobed, Figure 1e). Time lapse imaging revealed that abnormal acini containing polylobed cells do not show a development dependent decline of rotation speed (Supplementary Movie 1). Impaired acinus rotation was previously reported to prevent proper acinus development supporting our finding (Wang et al., 2013). By following the development of normal and abnormal acini over seven days, we quantified their relative size, hollowness and polarization over time (Figure 1f). Abnormal acini of *ESPL1* and *TOP2A* were smaller and showed a reduction in hollowness and polarization. Topoisomerase A (TOP2A) plays an important role in chromatin condensation and sister chromatid separation to relief torsional stress on DNA (Chen et al., 2015), similarly separase (ESPL1) cleaves cohesin subunits of sister chromatids to release progress to the mitosis machinery (Papi et al., 2005). The downregulation of topoisomerase A or separase lead to segregation defects affecting proper cytokinesis after mitosis (Figure 1g). The resulting polylobed cells hamper the proliferation and growth organization of MCF10A spheroids.

### Abnormal acini show a deregulation of cancer relevant genes

We examined the expression of *scrambled* (18 genes upregulated) and compared it to *ESPL1* and *TOP2A* with 67 and 283 upregulated genes (more than 1.5 -fold, p-value < 0.05), respectively. Gene enrichment of *ESPL1* upregulated genes indicated an association to breast cancer gene sets, whereas *TOP2A* showed a stronger association to leukemia gene sets (Supplementary Table 2). *Scrambled* was not enriched for any cancer related gene sets. The expression profiles contained a broad range of cancer related genes between *ESPL1* and *TOP2A* (52 genes, Figure 2a). These 52 genes were enriched for cytoskeleton, adhesion, EGFR pathway, breast cancer, and cancer module gene sets (Supplementary Table 2). Besides ALDH1A3 both profiles included CCND1 to be upregulated. Metabolic aldehyde dehydrogenase 1A3 and cyclin D1 are frequently deregulated in a variety of entities including breast cancer (Marcato et al., 2011, Musgrove et al., 2011). Immunostainings as well as qPCR confirmed that ALDH1A3 and CCND1 are upregulated in both aspects (Figure 2b, c). Since CCND1 plays an important role in cell cycle progression, we further questioned the mechanism of upregulation for CCND1. TCGA data suggest that CCND1 expression is insensitive against copy number variations (Supplementary Figure 1) and is rarely mutated or differentially methylated in invasive breast cancer (Tate et al., 2018). However, it is upregulated in nearly 20 % of invasive breast cancers (Tate et al., 2018) and plays a role in many other cancers (Musgrove et al., 2011). This led us to study whether disrupted chromatin interactions would play a role in CCND1 deregulation in aneuploid cells.

**Figure 2:**
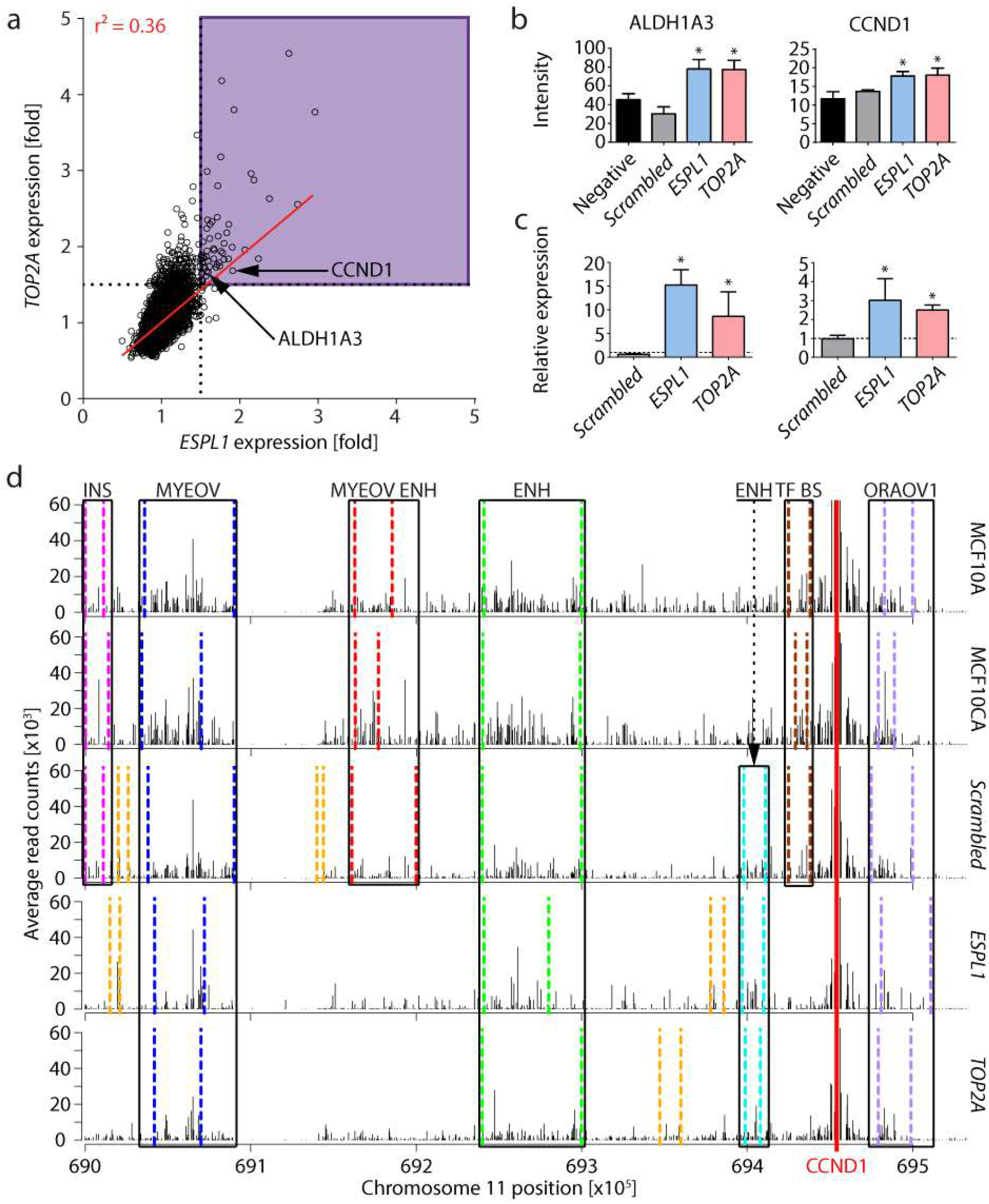
Aneuploid MCF10A acini show an altered expression of CCND1 as well as disrupted CCND1-promotor interactions. a) Correlation of the expression profiles of ESPL1 and TOP2A. b) Immunostaining of ALDH1A3 and CCND1 comparing MCF10A, *scrambled*, *ESPL1*, and *TOP2A*. c) Relative expression of CCND1 in *scrambled*, *ESPL1* and *TOP2A*. d) Interactions of the CCND1 promoter with surrounding regions identified by read counts calculated by PeakC (Geeven et al., 2018) for MCF10A, MCF10CA, *scrambled*, *ESPL1*, and *TOP2A*. Interactions that are common for more than one condition are boxed-in in black. Individual interactions in regions with low read coverage are highlighted in orange. Further explanations and references about regulatory elements in main text.

### Additional chromosomes interfere with chromatin interactions

We conducted chromosome conformation capture (4C) to test if the upregulation of CCND1 can be a result of disrupted interactions of the promoter and its regulatory regions in the context of aneuploidy. We focused on the near-cis region that contains a high read coverage on chromosome 11. The analysis of the interactions in proximity (window of approximately 1 MB around the CCND1 viewpoint) to the promoter revealed interactions conserved for all five conditions which are located at (i) the MYEOV promoter region, which was reported to be co-expressed with CCND1 in esophageal squamous cell carcinoma (Janssen et al., 2002, Figure 2d, blue), (ii) an enhancer/transcription factor binding (E/TFB) site (Figure 2d, green), and (iii) the adjacent ORAOV1 promoter region (11q13-associated driver of cancer, Zhai et al., 2014, Figure 2d, purple). Conversely, *ESPL1* and *TOP2A* did not show interactions with an insulator region (Figure 2d, pink), an MYEOV enhancer site (Figure 2d, red) and another transcription factor binding site (Figure 2d, brown). These positions were present for *scrambled*, MCF10A and MCF10CA (Figure 2d). In the region of another E/TFB site interactions could be detected for *ESPL1*, *TOP2A* and *scrambled* (Figure 2d, cyan). In conclusion, *ESPL1* and *TOP2A* share a specific regulatory interaction profile supposedly disturbed in comparison to normal MCF10 spheroids by chromosomal rearrangements or additional interactions.

## DISCUSSION

Early stages of breast cancer show comparable rates of aneuploidy to later stages (Ottesen et al., 1995, wang et al., 2014). This indicates not only an early occurrence of aneuploidy, but acquiring this aneuploidy might be an important step towards tumorigenesis. In the present work we try to mimic the early acquisition of aneuploidy found in breast tumorigenesis using 3D cultured MCF10A cells and study the effects on the morphogenesis and expression. Therefore, we utilize siRNA targeted against genes associated with aneuploidy and breast cancer.

The induction of aneuploidy in MCF10A cells by downregulation of ESPL1 and TOP2A lead to the formation of polylobed nuclei that frequently show mitotic abnormalities which are also regularly seen in cancer (Thompson, Bakhoum, & Compton, 2010). Additionally, the morphological resemblance of abnormal acini to lobular neoplasia and the abnormal expression of cancer related gene sets seem to make this an appropriate model to study early stages of breast cancer development. This is supported by the correlation of the expression profiles of MCF10A acini to breast cancer gene sets which was not measured for monolayer MCF10A cells.

Previous studies have already reported that aneuploidy alone is enough to induce the upregulation of cancer related genes (Zhang et al., 2013; Ben-david et al., 2014). This raises the question, why specific genes can be upregulated in such a seemingly random process. In our work we detected the upregulation of several pathways. The mitotic errors caused by ESPL1 and TOP2A downregulation results in the mis-segregation of whole chromosome sets resulting in near tetraploid karyotypes with some additional deletions or duplications of single chromosomes. How can the deregulation be accounted for by copy number changes of single chromosomes or genes only? Apparently, changes in number of single chromosomes can alter the expression profile completely as supported by other studies (Nawata et al., 2011, Kaushal et al., 2003).

*ESPL1* and *TOP2A* show a remarkably similar deregulation of specific genes as well as morphologies. Transcription factories are tightly controlled islands in the nucleus playing a central role in gene expression and genome organization. The mechanism of formation of transcription factories is not clear, yet (Marenduzzo et al., 2006, Schoenfelder et al., 2010, Cook & Marenduzzo, 2018). As seen from the images of polylobed nuclei, not only the chromatin content, but also the structure changes during interphase as well as mitosis for *ESPL1* and *TOP2A*. Given that the organization of chromatin interactions is tightly controlled and dependent on a correct chromosomal composition the accumulation of additional genetic material, potentially directly interferes with proper chromatin-chromatin interactions and could thus deregulate genes.

An aneuploidy dependent disruption of chromatin interactions might explain why for some genes copy number and expression do not correlate. In this concept, additional as well as missing chromosomes could disproportionally change expression levels through altered promoter-enhancer interactions overcoming sheer copy numbers. In the expression profiles of *ESPL1* and *TOP2A* we could find several genes that share transcription factor binding motifs of TCF3, PAX4, ETS2 and MAZ. Taking this into account it remains to be tested if the expression of genes with specific enhancer landscapes might be more susceptible against chromatin changes. This would be in line with the explanation that transcription factories form by accumulation of genes with similar enhancer requirements (Schoenfelder et al., 2010).

This work shows that aneuploidy can influence cancer development by altering the chromatin interaction landscape and subsequently the expression of cancer relevant genes. Although the mechanism is elusive the aneuploidization leads to the specific upregulation of CCND1 and formation of acini reminiscent of lobular neoplasia.

## MATERIAL AND METHODS

### Cell culture

MCF10A cells (Soule et al., 1990, ATCC, CRL-10317) transfected with a pBabe vector containing a construct of GFP labelled H2B were kindly provided the Zev Gartner and colleagues. MCF10AT (Dawson et al., 1996) and MCF10CA (Santner et al., 2001) were acquired from the Karmanos cancer institute (KCI, http://www.karmanos.org/home). The cells were cultured in polystyrene culture flasks (Greiner Bio-One GmbH, Kremsmünster, Austria, #690175) with DMEM/F12 medium (life technologies, ThermoFisher, Waltham, Massachusetts, USA, #11330-032) supplemented with 5 % serum (life technologies, ThermoFisher, Waltham, Massachusetts, USA, #1605-122), 10 μg/ml insulin (Sigma, St. Louis, Missouri, USA, I1882), 20 ng/ml EGF (Sigma, St. Louis, Missouri, USA, E9644), 500 μg/ml hydrocortisone (Sigma, St. Louis, Missouri, USA, H0888) and 100 ng/ml cholera toxin (Sigma, St. Louis, Missouri, USA, C8052). The cells were detached using 0.05 % trypsin/EDTA and propagated every 3-4 days.

To culture MCF10A cells as 3D culture 70 μl Matrigel® (Corning, New York, USA, #356231) was added to each well of a 24-well plate. After solidification of the layer 15.000 cells were seeded into each well. The medium was exchanged every 3 days.

### SiRNA mediated knockdown

SiRNA (Ambion, ThermoFisher, Waltham, Massachusetts, USA) transfection was conducted after a slightly modified supplier protocol. 1 μl Lipofectamine 2000 (ThermoFisher, Waltham, Massachusetts, USA, #11668019) was incubated in 50 μl nuclease free water (NFW) for 5 minutes at RT and subsequently mixed with 3 pmol siRNA in 50 μl NFW. The transfection mix was added to the cells that were washed with PBS once and then incubated for >6 hours at 37 °C. Afterwards the medium was exchanged and cells were further processed from day 2 after transfection.

### RNA extraction

RNA was extracted using the QIAGEN RNeasy extraction kit (Qiagen, Venlo, Netherlands, # 74106) after supplier protocol. The cells were lysed in the well using 350 μl RLT buffer. After 1-minute incubation at RT the lysate was mixed with 350 μl of 70 % ethanol. Afterwards the mixture was loaded onto RNeasy Mini spin columns, centrifuged at 8000g, and washed with RW1 and 2x RPE buffers successively. The RNA was eluted using NFW provided in the kit.

### QPCR quantification

The total RNA was treated with the Fermentas DNase digestion kit (ThermoFisher, Waltham, Massachusetts, USA, #K1622) after supplier recommendations to get rid of remaining DNA. In the next step the RNA was used for cDNA synthesis with the Fermentas RevertAID cDNA synthesis kit (ThermoFisher, Waltham, Massachusetts, USA, #K1622) after supplier recommendations. The cDNA was then diluted with NFW to an endconcentration of 1.4 ng/μl. 4 μl of diluted cDNA was mixed with 10 μl TaqMan Universal Master Mix II, with UNG (ThermoFisher, Waltham, Massachusetts, USA, # 4440044) and 1 μl TaqMan Gene Expression Assay (TermoFisher, Waltham, Massachusetts, USA, #4331182) with <100 ng cDNA and 5μl water in the designated well of a 96-well Q-PCR plate. The plate was measured and analyzed using the Fischer StepOnePlus RealTime PCR machine (ThermoFisher, Waltham, Massachusetts, USA, # 4376599) and the Step One Software version 2.0. The experiment was set to quantitation-comparative Ct (ΔΔCt), 6-FAM dye reagents with 95 °C 10 minutes denaturation, 40 cycles of 15 seconds at 95 °C, 30 seconds at 60 °C, melt curve: 95 °C for 15 seconds, 60 °C for 1 minute and a subsequent heating ramp of 0.3 °C/minute until 95 °C was reached. In the analysis RPLP0 served as housekeeping gene.

### Immunostaining

The cells were fixed using 2 % formaldehyde (Sigma, St. Louis, Missouri, USA, F8775) in PBS for 12 minutes at RT and subsequently washed 3x with PBS for 5 minutes at RT. Permeabilization was done with 0.5 % Triton X-100 for 10 minutes at 4 °C. After washing the cells 3x for 5 minutes at RT blocking solution (10 % serum + washing buffer: 38 g NaCL, 9.38 g Na2HPO4, 2.07 g NaH2PO4, 2.5 g NaN3, 5 g BSA, 10 ml Triton-X 100, 2.05 ml Tween-20, filled up to 500 ml with water and titrated to a pH of 7.4) was applied and incubated for 1 hour at RT. The blocking solution was then replaced with primary antibodies in blocking solution and incubated at 4 °C overnight. The next day the cells were washed twice with washing buffer for 5 minutes at RT each and then incubated with the secondary antibodies in blocking solution for 1 hour at RT. Afterwards the cells were washed once with washing buffer and once with PBS and subsequently imaged.

### Image settings for high content screen and time lapses

If not otherwise stated confocal imaging was conducted using the Zeiss LSM 780 (EC Plan-Neofluar 10x/0.30 M27) under the control of AutofocusScreen macro (“www.ellenberg.embl.de/apps/AFS”, 24.02.2016). A grid with nine positions per well was imaged.

Time lapse images were acquired using the Plan-Apochromat 20x/0.80 M27 objective with a digital zoom of 1.6x with an interval of 20 minutes and 17 stacks á 4 μm over 6 days. Two random positions per well were imaged.

### Image analysis and tracking

All images were analyzed using custom workflows within KNIME (Berthold et al., 2007). The high content screening data was calculated as the average per spheroid (3D) or field of view (2D) per condition for each parameter (spheroid count, spheroid size [suqare pixels], roundness, hollowness [H2B rim intensity / H2B core intensity], polarization [GM130 core intensity / GM130 rim intensity], Tamura texture, Haralick texture, H2AFX intensity, H2B intensity). The clusters were calculated using PCA and subsequent k-means clustering.

Time lapse images were automatically cropped into single spheroid 3D datasets. These 3D datasets were analyzed using 3D segmentation for the whole spheroids. Size, polarization, and hollowness were then calculated for each time point per single spheroid. To track single nuclei in these spheroids a Z-projection was analyzed using ImageJ/FIJIs Trackmate tracker within the KNIME workflow.

### Chromosome spreads

Cells with a density of 40-60 % (2D) or 6 days after seeding onto Matrigel (3D) were treated with 40 ng/ml colcemid (Sigma-Aldrich, St. Louis, Missouri, Vereinigte Staaten, D7385) for 20 hours at 37 °C. In the next step the cells were detached by adding 0.05 % T/E for 10 minutes at 37 °C (2D) or removed from Matrigel using the BD recovery solution after supplier recommendations and dissociated using 0.25 % T/E for 20 minutes at 37 °C and transferred into a reaction tube. The cells were then centrifuged at 300 g for 10 minutes at RT. The supernatant was aspirated until there was ~500 μl left and the cells resuspended by flicking the tube. Subsequently the cells were treated with 5 ml hypotonic solution (0.55 % KCl in deionized water) which was applied dropwise for 10 minutes at RT. Afterwards the cells were centrifuged at 300 g for 10 minutes at RT and the supernatant again aspirated until there was ~500 μl left so that the cells could be resuspended by flicking the tube. The cells in the suspension were then fixed by dropwise adding 5 ml of Carnoy’s Fixative (75 % Methanol, 25% acetic acid) and incubated for 10 minutes at RT. The fixation was repeated twice. If not otherwise stated the M-FISH on metaphase chromosome spreads (MCS) were prepared after Geigl et al., 2006 by the human genetics working group of Anna Jauch. DOP-PCR amplified probe pools were combinatorially labeled (DEAC, FITC, Cy3, Cy3.5, Cy5, Cy5.5, and Cy7) and hybridized with cot-1 DNA. Images of 15 MCSs were acquired using the LEICA DM RXA epifluorescence microscope (Leica Microsystems, Bensheim, Germany). The microscope was under control of the Leica Q-FISH software. The images were processed using the Leica MCK software (Leica Microsystems Imaging solutions, Cambridge, United Kingdom).

For the quantification of several MCSs the probes were further processed in collaboration with Metasystems and acquired as well as analyzed using the Metafer 4 v3.11.7 software (Metasystems, Altlußheim, germany).

### Microarray expression profiles and GSEA enrichment analysis

Extracted RNA was further processed by the DKFZ genomics core facility and hybridized onto Illumina HumanHT-12 v4 Expression BeadChips and quantile normalized. Significantly deregulated genes (1.5-fold, p<=0.05) were overlapped with all relevant gene sets of the molecular signature database GSEA (Subramanian et al., 2005).

### Chromosome conformation capture (4C)

We extracted 10 million cells per viewpoint from Matrigel using dispase. Subsequently, the cells were dissociated in 0.05 % trypsin/EDTA and further processed as previously described (Van de Werken et al., 2012). We used DPNII or NIaIII as primary and Csp6I or Bfal as secondary restriction enzymes. The primer sequences with the secondary cutter (BfaI in the case of CCND1) were used to demultiplex the reads. In silico digestion of grch37/hg19 reference genome with both primary and secondary cut enzymes were performed using FourCSeq (Klein et al., 2015). The clipped reads were mapped to grch37/hg19 whole genome sequences. Only the reads starting at the cutting site of the second cutter were retained for the following analysis. The normalization between samples was performed according to the library size. To estimate the genomic range of the near-cis interactions we used 4C-ker (Raviram et al., 2016). The read counts were analyzed using the R package PeakC (Geeven et al., 2018) with a flanking window size of 700kb around the viewpoint, a running window size of 10 and false-discovery ratio threshold of 0.01. The definition of genomic regions was done using the UCSC Genome Browser (Kent et al., 2002) and ENCODE annotation data (Rosenbloom et al., 2013). Datasets with low quality were identified using the ratio of reads in cis and trans with a cutoff of 75% and subsequently excluded.

### Statistics

The data quantified by KNIME custom workflows was further processed by integrated custom R scripts and statistically quantified using linear regression, linear/non-linear fitting or ANOVA where appropriate utilizing the functions of Graphpad v 6.07. A statistical significance was considered to be at p ≤ 0.05.

## ACKNOWLEDGEMENT

We thank Holger Erfle und Jürgen Behnke (BioQuant Center Heidelberg University) for technical laboratory support and Elzo de Wit (Netherlands Cancer Institute) for support and helpful discussions regarding 4C analysis. We thank the microarray unit of the DKFZ Genomics and Proteomics Core Facility for providing the Illumina Whole-Genome Expression Beadchips and related services.

## FUNDING

This study was supported by the BMBF-funded Heidelberg Center for Human Bioinformatics (HD-HuB) within the German Network for Bioinformatics Infrastructure (de.NBI) (#031A537A, # 031A537C) and DKFZ-HIPO through Grant No. HIPO-H012.

## CONTRIBUTIONS

MW, CC and RE conceived the study, MW and CC designed experiments; MW performed cell culture experiments, siRNA treatments, RNA extraction, QPCR experiments, immunofluorescence stainings, confocal microscopy, chromosome preparation and 4C data analysis; SA and KJ have conducted DNA preparation and 4C experiments; QW, PS and CH have processed and quality checked 4C data; CK and MS have conducted mouse experiments; LM, CD and MW have developed image processing and analysis workflows; ST has developed acinus preparation for 4C experiments; BS and AJ have conducted and analyzed 24 multicolor FISH experiments; IC has quantified metaphase spreads; DD has developed 4C experiment workflows; MW, CC and RE wrote the manuscript. All authors revised and approved the manuscript.

## SUPPLEMENTARY DATA

**Supplementary Figure 1:**
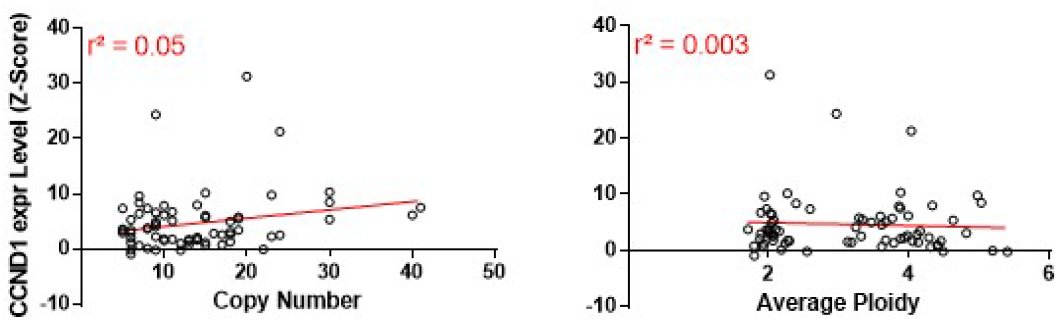
Correlation of CCND1 allele copy number or average ploidy with CCND1 expression level in invasive breast cancer according to TCGA data.

**Supplementary Table 1:** List of all screening candidates and their corresponding reference. The gene names and IDs are according to NCBI.

| Gene | ID | Reference |
| --- | --- | --- |
| ACTA1 | 58 | Havaki 2007 |
| APC | 324 | Thompson et al., 2010, Tate et al., 2018 |
| ARID1A | 8289 | Tate et al., 2018 |
| ATM | 472 | Pandita 2002, Tate et al., 2018 |
| ATR | 545 | Pino & Chung 2010 |
| AURKB | 9212 | Carter et al., 2006 |
| BAZ2A | 11176 | Parry and Clarke 2011 |
| BAZ2B | 29994 | Parry and Clarke 2011 |
| BRCA1 | 672 | Thompson et al., 2010, Tate et al., 2018 |
| BRCA2 | 675 | Thompson et al., 2010 |
| C20orf24 | 55969 | Carter et al., 2006 |
| CAMLG | 819 | Thompson et al., 2010 |
| CDC45 | 8318 | Chibon et al., 2010 |
| CDC6 | 990 | Carter et al., 2006 |
| CDH1 | 999 | Tate et al., 2018 |
| CDKN1B | 1027 | Davoli et al., 2013 |
| CDKN2A | 1029 | Davoli et al., 2013 cell, Tate et al., 2018 |
| CENPE | 1062 | Chibon et al., 2010 |
| CENPF | 1063 | Thompson et al., 2010 |
| CHEK1 | 1111 | Carter et al., 2006 |
| CHFR | 55743 | Perez de Castro et al., 2007 |
| CKAP2 | 26586 | Thompson et al., 2010 |
| CMAS | 55907 | Carter et al., 2006 |
| COPB2 | 9276 | Hutchins et al., 2010 |
| CTPS1 | 1503 | Carter et al., 2006 |
| DIAPH3 | 81624 | Johansson 2013 |
| ECT2 | 1894 | Carter et al., 2006 |
| ESPL1 | 9700 | Carter et al., 2006 |
| FBXW7 | 55294 | Thompson et al., 2010 |
| FHIT | 2272 | Saldivar 2012 |
| FOXM1 | 2305 | Carter et al., 2006 |
| GATA3 | 2625 | Tate et al., 2018 |
| H2AFX | 3014 | Carter et al., 2006 |
| HAUS8 | 93323 | Thompson et al., 2010 |
| INCENP | 3619 | Neumann et al., 2010 |
| KIF2B | 84643 | Thompson et al., 2010 |
| KIF4A | 24137 | Carter et al., 2006 |
| KMT2C | 58508 | Tate et al., 2018 |
| LATS1 | 9113 | Sorrentino et al., 2014 |
| LATS2 | 26524 | Sorrentino et al., 2014 |
| LMNB2 | 84823 | Kuga et al., 2014 |
| LLGL1 | 3996 | Russ et al., 2012 |
| LLGL2 | 3993 | Russ et al., 2012 |
| MAD2L1 | 4085 | Carter et al., 2006 |
| MAP2K4 | 6416 | Carter et al., 2006, Tate et al., 2018 |
| MAP3K1 | 4214 | Davoli et al., 2013 |
| MAPRE1 | 22919 | Thompson et al., 2010 |
| MBD5 | 55777 | Parry and Clarke 2011 |
| MBD6 | 114785 | Parry and Clarke 2011 |
| MYH10 | 4628 | Overholzer 2007 |
| MYO10 | 4651 | Overholzer 2007 |
| NCAPH | 23397 | Chibon et al., 2010 |
| NEU1 | 4758 | Butler et al., 2013 |
| NISCH | 11188 | Johansson et al., 2013 |
| NUP98 | 4928 | Perez de Castro et al., 2007 |
| PARD3 | 56288 | Wan et al., 2013 |
| PLK4 | 10733 | Chibon et al., 2010 |
| PRC1 | 9055 | Carter et al., 2006 |
| PTEN | 5728 | Tate et al., 2018 |
| RANBP1 | 5902 | Thompson et al., 2010 |
| RB1 | 5925 | Thompson et al., 2010, Tate et al., 2018 |
| RBMX | 27316 | Davoli et al., 2013 |
| RHOA | 387 | Overholzer 2007 cell |
| ROCK1 | 6093 | Overholzer 2007 cell |
| ROCK2 | 9475 | Overholzer 2007 cell |
| SETDB2 | 83852 | Parry and Clarke 2011 |
| SGO2 | 151246 | Chibon et al., 2010 |
| SMAD2 | 4087 | Petersen et al., 2010 |
| SPINK7 | 84651 | Thompson et al., 2010 |
| TOP2A | 7153 | Carter et al., 2006 |
| TP53 | 7157 | Thompson et al., 2010, Tate et al., 2018 |
| TP63 | 8626 | Stefanou et al., 2004 |
| TPX2 | 22974 | Carter et al., 2006 |
| TRIP13 | 9319 | Carter et al., 2006 |
| TTN | 7273 | Tate et al., 2018 |
| UHRF2 | 115426 | Parry and Clarke 2011 |
| VCL | 7414 | Mierke et al., 2010 |
| ZBTB4 | 57659 | Parry and Clarke 2011 |
| ZFP36L1 | 677 | Davoli et al., 2013 |
| ZSCAN22 | 342945 | Personal communication (Claudia Lukas, Uni Kopenhagen) |
| ZWILCH | 55055 | Perez de Castro et al., 2007 |
| ZWINT | 11130 | Carter et al., 2006 |

**Supplementary Table 2:**
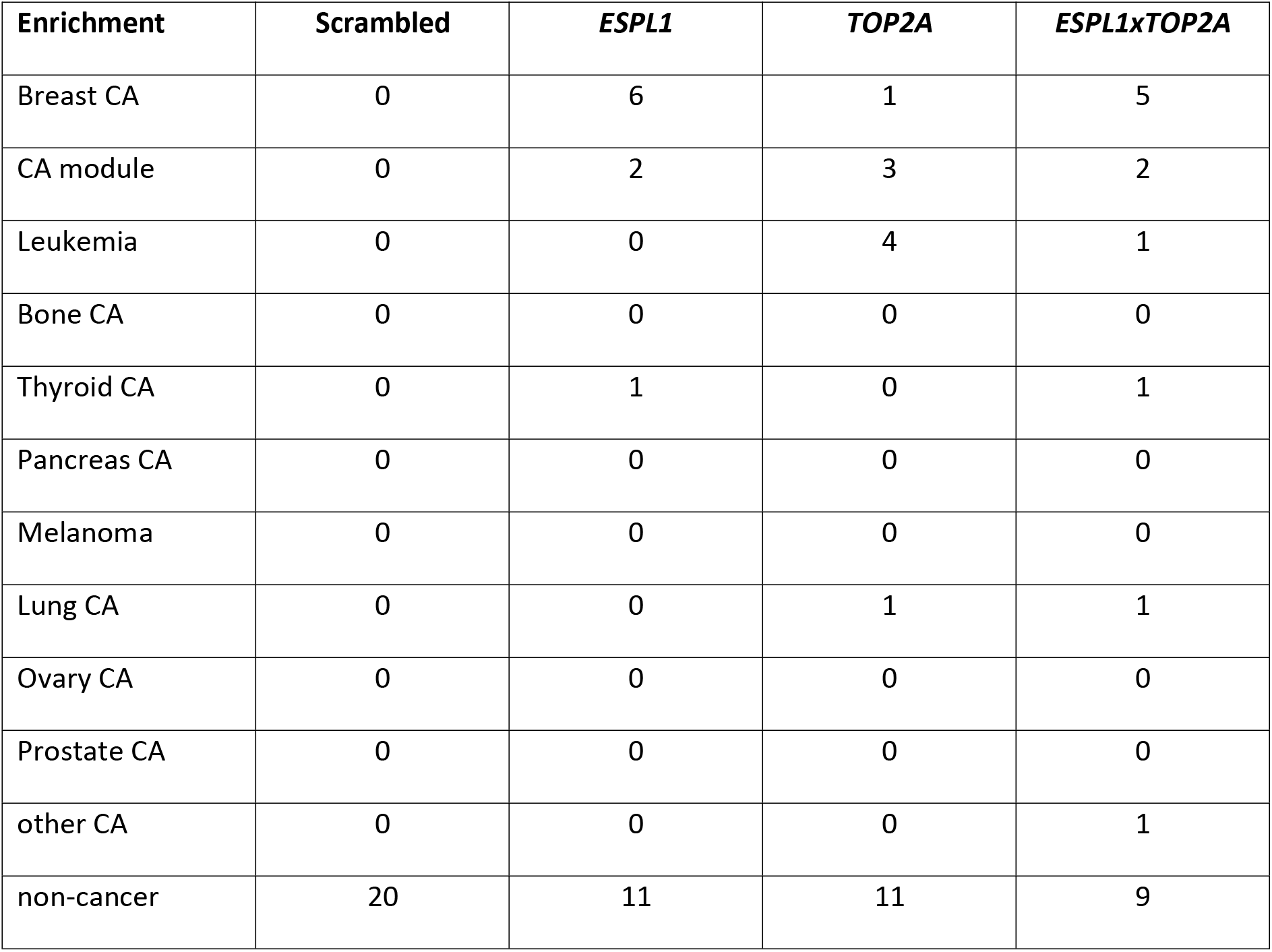
Top 20 occurrences of genes sets associated to different cancer entities after gene set enrichment analysis (Subramanian et al., 2005) of upregulated genes of *scrambled*, *ESPL1*, *TOP2A*, and the consensus of upregulated genes of *ESPL1* and *TOP2A* (*ESPL1xTOP2A*). CA: Cancer.

| Enrichment | Scrambled | *ESPL1* | *TOP2A* | *ESPL1xTOP2A* |
| --- | --- | --- | --- | --- |
| Breast CA | 0 | 6 | 1 | 5 |
| CA module | 0 | 2 | 3 | 2 |
| Leukemia | 0 | 0 | 4 | 1 |
| Bone CA | 0 | 0 | 0 | 0 |
| Thyroid CA | 0 | 1 | 0 | 1 |
| Pancreas CA | 0 | 0 | 0 | 0 |
| Melanoma | 0 | 0 | 0 | 0 |
| Lung CA | 0 | 0 | 1 | 1 |
| Ovary CA | 0 | 0 | 0 | 0 |
| Prostate CA | 0 | 0 | 0 | 0 |
| other CA | 0 | 0 | 0 | 1 |
| non-cancer | 20 | 11 | 11 | 9 |

**Supplementary Movie 1:**
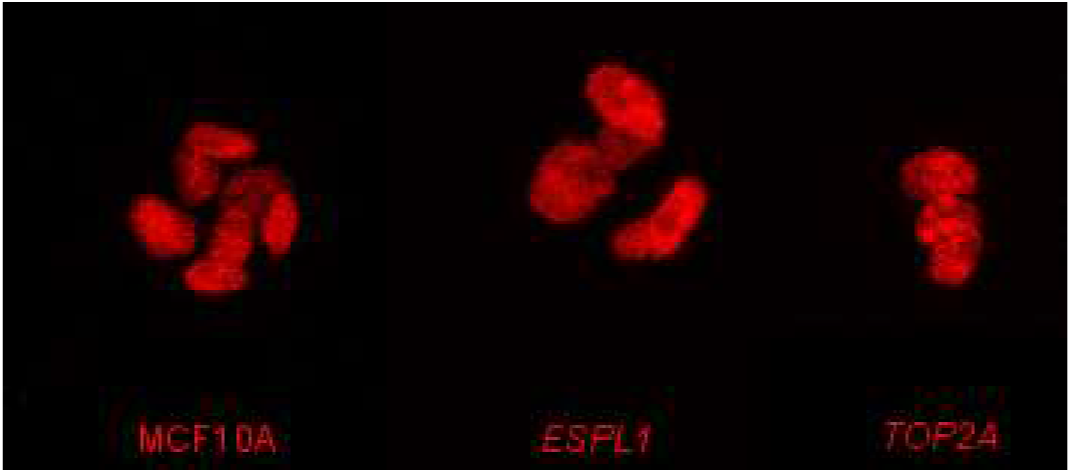
Acini of MCF10A, *ESPL1*, and *TOP2A* (*from left to right*) over 276 time points (6 days, 30-minute interval).

## REFERENCES

Ben-David U., Arad G., Weissbein U., Mandefro B., Maimon A., Golan-Lev T., Narwani K., Clark AT., Andrews P.W., Benvenisty N., Carlos Biancotti J., 2014. Aneuploidy induces profound changes in gene expression, proliferation and tumorigenicity of human pluripotent stem cells. Nature Communications, 5:1–11.

Berthold M.R., Cebron N., Dill F., Gabriel T.R., Koetter T., Meinl T., Ohl P., Sieb C., Thiel K., Wiswedel B., 2007. KNIME – The Konstanz Information Miner. Springer, 11:26–31.

Boveri T., 2008. Concerning the origin of malignant tumours by Theodor Boveri. Translated and annotated by Henry Harris. J Cell Sci., 121:1–84.

Bray F., Ferlay J., Soerjomataram I., Siegel R.L., Torre L.A., Jemal A., 2018. Global cancer statistics 2018: GLOBOCAN estimates of incidence and mortality worldwide for 36 cancers in 185 countries. CA Cancer J Clin. 68:394–424.

Carter S.L., Eklund A.C., Kohane I.S., Harris L.N., Szallasi Z., 2006. A signature of chromosomal instability inferred from gene expression profiles predicts clinical outcome in multiple human cancers. Nature genetics, 38:1043–1048.

Chen T., Sun Y., Ji P., Kopetz S., Zhang W., 2015. Topoisomerase IIα in chromosome instability and personalized cancer therapy. Oncogene, 34:4019–31.

Cimini, D., 2008. Merotelic kinetochore orientation, aneuploidy, and cancer. Biochimica et Biophysica Acta - Reviews on Cancer, 1786, pp.32–40.

Cook P.R., 2009. A model for all genomes: the role of transcription factories. J Mol Biol. 395:1–10.

Cook P.R., Marenduzzo D., 2018. Transcription-driven genome organization: a model for chromosome structure and the regulation of gene expression tested through simulations. Nucleic Acids Res. 46:9895–9906.

Dawson P.J., Wolman S.R., Tait L., Heppner G.H., Miller F.R., 1996. MCF10AT: a model for the evolution of cancer from proliferative breast disease. Am J Pathol. 148:313–9.

Debnath J., Muthuswamy S.K., Brugge J.S., 2003. Morphogenesis and oncogenesis of MCF-10A mammary epithelial acini grown in three-dimensional basement membrane cultures. Methods, 30:256–268.

Dekker J., Rippe K., Dekker M., Kleckner N., 2002. Capturing Chromosome Conformation. Science, 295:1306–1311.

Gao R., Davis A., McDonald T.O., Sei E., Shi X., Wang Y., Tsai P.C., Casasent A., Waters J., Zhang H., Meric-Bernstam F., Michor F, Navin N.E., 2016. Articles Punctuated copy number evolution and clonal stasis in triple-negative breast cancer. Nature genetics, 48:1119–1130.

Geeven G., Teunissen H., de Laat W., de Wit E., 2018. peakC: a flexible, non-parametric peak calling package for 4C and Capture-C data. Nucleic Acids Res. 46:e91.

Geiger T., Cox J., Mann M., 2010. Proteomic Changes Resulting from Gene Copy Number Variations in Cancer Cells. PLoS Genet. 6:e1001090.

Geigl, J.B., Uhrig, S., Speicher, M.R., 2006. Multiplex-fluorescence in situ hybridization for chromosome karyotyping. Nature protocols, 1:1172–1184.

Janssen J.W., Imoto I., Inoue J., Shimada Y., Ueda M., Imamura M., Bartram C.R., Inazawa J., 2002. MYEOV, a gene at 11q13, is coamplified with CCND1, but epigenetically inactivated in a subset of esophageal squamous cell carcinomas. J Hum Genet. 47:460–4.

Kaushal D., Contos J.J., Treuner K., Yang A.H., Kingsbury M.A., Rehen S.K., McConnell M.J., Okabe M., Barlow C., Chun J., 2003. Alteration of gene expression by chromosome loss in the postnatal mouse brain. J Neurosci. 23:5599–606.

Kent W.J., Sugnet C.W., Furey T.S., Roskin K.M., Pringle T.H., Zahler A.M., Haussler D., 2002. The human genome browser at UCSC. Genome Res. 12:996–1006.

Klein F.A., Pakozdi T., Anders S., Ghavi-Helm Y., Furlong E.E., Huber W., 2015. FourCSeq : analysis of 4C sequencing data. Bioinformatics, 31:3085–3091.

Lieberman-Aiden E., van Berkum N.L., Williams L., Imakaev M., Ragoczy T., Telling A., Amit I., Lajoie B.R., Sabo P.J., Dorschner M.O., Sandstrom R., Bernstein B., Bender M.A., Groudine M., Gnirke A., Stamatoyannopoulos J., Mirny L.A., Lander E.S., Dekker J., 2009. Comprehensive Mapping of Long-Range Interactions Reveals Folding Principles of the Human Genome. Science, 326:289–294.

Lundberg A., Lindström L.S., Li J., Harrell J.C., Darai-Ramqvist E., Sifakis E.G., Foukakis T., Perou C.M., Czene K., Bergh J., Tobin N.P., 2019. The long-term prognostic and predictive capacity of cyclin D1 gene amplification in 2305 breast tumours. Breast Cancer Res. 21: 34.

Marcato P., Dean C.A., Pan D., Araslanova R., Gillis M., Joshi M., Helyer L., Pan L., Leidal A., Gujar S., Giacomantonio C.A., Lee P.W., 2011. Aldehyde Dehydrogenase Activity of Breast Cancer Stem Cells is Primarily Due to Isoform ALDH1A3 and Its Expression is Predictive of Metastasis. STEM CELLS, 29:32–45.

Marenduzzo D., Micheletti C., Cook P.R., 2006.Entropy-driven genome organization. Biophys J. 90:3712–21.

Miele A., Dekker J., 2009. Mapping cis- and trans- chromatin interaction networks using chromosome conformation capture (3C). Methods Mol Biol. 464:105–21.

Musgrove E.A., Caldon C.E., Barraclough J., Stone A., Sutherland R.L., 2011. Cyclin D as a therapeutic target in cancer. Nature reviews. Cancer, 11:558–572.

Nawata H., Kashino G., Tano K., Daino K., Shimada Y., Kugoh H., Oshimura M., Watanabe M., 2011. Dysregulation of gene expression in the artificial human trisomy cells of chromosome 8 associated with transformed cell phenotypes. PLoS One. 6:e25319.

Nowak M.A., Komarova N.L., Sengupta A., Jallepalli P.V., Shih IeM., Vogelstein B., Lengauer C., 2002. The role of chromosomal instability in tumor initiation. Proceedings of the National Academy of Sciences, 99:16226–16231.

Osborne C.S., Chakalova L., Brown K.E., Carter D., Horton A., Debrand E., Goyenechea B., Mitchell J.A., Lopes S., Reik W., Fraser P., 2004. Active genes dynamically colocalize to shared sites of ongoing transcription. Nat Genet. 36:1065–71.

Ottesen G.L., Christensen I.J., Larsen J.K., Kerndrup G.B., Hansen B., Andersen J.A., 1995. DNA aneuploidy in early breast cancer. British journal of cancer, 72:832–9.

Otto, T. & Sicinski, P., 2017. Cell cycle proteins as promising targets in cancer therapy. Nature reviews. Cancer, 17:93–115.

Papi M., Berdougo E., Randall C.L., Ganguly S., Jallepalli P.V., 2005. Multiple roles for separase auto-cleavage during the G2 / M transition. Nature Cell Biology, 7:1029–1035.

Raviram R., Rocha P.P., Müller C.L., Miraldi E.R., Badri S., Fu Y., Swanzey E., Proudhon C., Snetkova V., Bonneau R., Skok J.A., 2016. 4C-ker: A Method to Reproducibly Identify Genome-Wide Interactions Captured by 4C-Seq Experiments. PLoS Comput Biol. 12e1004780.

Rosenbloom K.R., Sloan C.A., Malladi V.S., Dreszer T.R., Learned K., Kirkup V.M., Wong M.C., Maddren M., Fang R., Heitner S.G., Lee B.T., Barber G.P., Harte R.A., Diekhans M., Long J.C., Wilder S.P., Zweig A.S., Karolchik D., Kuhn R.M., Haussler D., Kent W.J., 2013. ENCODE data in the UCSC Genome Browser: year 5 update. Nucleic Acids Res. 41(Database issue):D56–63.

Santner S.J., Dawson P.J., Tait L., Soule H.D., Eliason J., Mohamed A.N., Wolman S.R., Heppner G.H, Miller F.R., 2001. Malignant MCF10CA1 cell lines derived from premalignant human breast epithelial MCF10AT cells. Breast cancer research and treatment, 65:101–10.

Schoenfelder S., Clay I., Fraser P., 2010. The transcriptional interactome: gene expression in 3D. Curr Opin Genet Dev. 20:127–33.

Simonis M., Klous P., Splinter E., Moshkin Y., Willemsen R., de Wit E., van Steensel B., de Laat W., 2006. Nuclear organization of active and inactive chromatin domains uncovered by chromosome conformation capture–on-chip (4C). Nature genetics, 38:1348–1354.

So J.Y., Lee H.J., Kramata P., Minden A., Suh N., 2012. Differential Expression of Key Signaling Proteins in MCF10 Cell Lines, a Human Breast Cancer Progression Model. Mol Cell Pharmacol, 4:31–40.

Sotillo R., Schvartzman J.M., Socci N.D., Benezra R., 2010. Mad2-induced chromosome instability leads to lung tumour relapse after oncogene withdrawal. Nature, 464:436–40.

Soule H.D., Maloney T.M., Wolman S.R., Peterson W.D. Jr., Brenz R., McGrath C.M., Russo J., Pauley R.J., Jones R.F., Brooks S.C., 1990. Isolation and characterization of a spontaneously immortalized human breast epithelial cell line, MCF-10. Cancer Res. 50:6075–86.

Subramanian A., Tamayo P., Mootha V.K., Mukherjee S., Ebert B.L., Gillette M.A., Paulovich A., Pomeroy S.L., Golub T.R., Lander E.S., Mesirov J.P., 2005. Gene set enrichment analysis: A knowledge-based approach for interpreting genome-wide expression profiles. Proc. Natl. Acad. Sci. 102, 15545–15550.

Tate J.G., Bamford S., Jubb H.C., Sondka Z., Beare D.M., Bindal N., Boutselakis H., Cole C.G., Creatore C., Dawson E., Fish P., Harsha B., Hathaway C., Jupe S.C., Kok C.Y., Noble K., Ponting L., Ramshaw C.C., Rye C.E., Speedy H.E., Stefancsik R., Thompson S.L., Wang S., Ward S., Campbell P.J., Forbes S.A., 2018. COSMIC: the Catalogue Of Somatic Mutations In Cancer. Nucleic Acids Res. 47:D941–D947.

Thompson S.L., Bakhoum S.F., Compton D.A., 2010. Mechanisms of chromosomal instability. Curr Biol. 20:R285–95.

Van de Werken H.J., Landan G., Holwerda S.J., Hoichman M., Klous P., Chachik R., Splinter E., Valdes-Quezada C., Oz Y., Bouwman B.A., Verstegen M.J., de Wit E., Tanay A., de Laat W., 2012. Robust 4C-seq data analysis to screen for regulatory DNA interactions. Nat Methods. 9:969–72.

Wang H., Lacoche S., Huang L., Xue B., Muthuswamy S.K., 2013. Rotational motion during three-dimensional morphogenesis of mammary epithelial acini relates to laminin matrix assembly. Proceedings of the National Academy of Sciences of the United States of America, 110:163–8.

Wang Y., Waters J., Leung M.L., Unruh A., Roh W., Shi X., Chen K., Scheet P., Vattathil S., Liang H., Multani A., Zhang H., Zhao R., Michor F., Meric-Bernstam F., Navin N.E., 2014. Clonal evolution in breast cancer revealed by single nucleus genome sequencing. Nature, 512:1–15.

Zhai C., Li Y., Mascarenhas C., Lin Q., Li K., Vyrides I., Grant C.M., Panaretou B., 2014. The function of ORAOV1/LTO1, a gene that is overexpressed frequently in cancer: essential roles in the function and biogenesis of the ribosome. Oncogene. 33:484–94.

Zhang R., Hao L., Wang L., Chen M., Li W., Li R., Yu J., Xiao J., Wu J., 2013. Gene expression analysis of induced pluripotent stem cells from aneuploid chromosomal syndromes. BMC Genomics, 14:8.

